# The double tubular contractile structure of the type VI secretion system displays striking flexibility and elasticity

**DOI:** 10.1101/470229

**Authors:** Maria Silvina Stietz, Xiaoye Liang, Megan Wong, Steven Hersch, Tao G. Dong

**Affiliations:** Ecosystem and Public Health; Snyder Institute for Chronic Diseases; Biochemistry and Molecular Biology, Cumming School of Medicine, University of Calgary, Calgary, AB, T2N4Z6, Canada

**Keywords:** type VI secretion system, bacteriophage, spheroplast, elastic rod, elasticity, contractile tubes

## Abstract

The double tubular structure of the type VI secretion system (T6SS) is considered as one of the longest straight and rigid intracellular structures in bacterial cells. Contraction of the T6SS outer sheath occurs almost instantly and releases sufficient power to inject the inner needle-like Hcp tube and its associated effectors into target bacterial cells through piercing the stiff cell envelope. The molecular mechanism triggering T6SS contraction remains elusive. Here we report that the double tubular T6SS structure is strikingly flexible and elastic, forming U-, circular-, or S-shapes while maintaining functional for contraction and substrate delivery. We show that physical contact with cytoplasmic membrane induced a range of T6SS structure deformation, but the resultant mechanical pressing force on the T6SS baseplate did not trigger contraction. Our results also reveal a stalling intermediate stage of sheath-tube extension following which the structure contracts or resumes to extend. These observations suggest that the recruitment equilibrium of sheath-tube precursors to the extending structure is key to stability/contraction and lead us to propose a model of T6SS contraction, termed ESCAPE (extension-stall-contraction and precursor equilibrium). Our data highlight the remarkable flexibility of the double tubular T6SS structure and its length control mechanism distinct from the other evolutionarily related contractile cell-puncturing systems.

## Introduction

Bacteria have evolved with various mechanisms to protect themselves and outcompete neighboring species in complex communities ^1^. The antagonistic interactions mediated by the activities of those bacterial weapons could dictate community structures and dynamic changes ^2–4^. The type VI secretion system (T6SS) is one such lethal weapon used by many Gram-negative bacteria to deliver toxic effectors into neighboring bacterial and eukaryotic cells^5–8^. Encoded by 13 conserved genes, the T6SS structure consists of a membrane complex TssJLM, a baseplate TssK-TssEFG, and a double tubular structure comprising an outer VipA/B (TssB/C) sheath encircling an inner needle-like Hcp tube ^9–15^. At the tip of the tube sits a spike complex VgrG/PAAR that not only sharpens the tube but also serves as a main loading site for effectors ^16–19^. The inner tube and the outer sheath share the same six-start helical structures with each ring composed of hexamers of Hcp and heterodimeric VipA/B, respectively ^13,14^. The T6SS structure assembles from the baseplate and polymerizes inward with new subunits added to the distal end, often spanning across the cell ^10^. The extended sheath is in a high energy conformation, and its contraction is proposed to initiate from the baseplate side to the distal end, shortening the sheath by half within two milliseconds ^10,20^. After contraction, the N-terminus of VipB is exposed and recognized by the AAA+ (<u>A</u>TPase <u>A</u>ssociated with various cellular <u>A</u>ctivities) ATPase ClpV that disassembles the contracted sheath and enables recycling of sheath subunits for another assembly ^21–24^. By contrast, the inner Hcp tube is ejected and requires *de novo* synthesis, hence under distinct transcriptional regulation from the sheath subunits ^25^.

Size determination of a cellular organelle is a critical control for its proper function, and yet it often remains poorly defined mechanistically due to the complexity ^26–28^. Molecular ruler proteins have been discovered for length control of the type III secretion system (T3SS) and its related flagellum hook that form extracellular structures protruding from the outer membrane ^29–31^, as well as for length control of phage tail ^32^. By contrast, it remains elusive regarding the mechanisms that dictate the intracellular T6SS sheath-tube length, terminate extension and trigger contraction. The T6SS extended sheath exhibits a quite narrow length distribution in normal rod cells ^10,33^, suggesting random contraction is rare and the size is somehow controlled. However, a specific ruler protein is likely absent because the extended sheath-tube structure size has been found to decrease when Hcp is limited and increase in spheroplast cells ^10,34^. Because the T6SS double tubular structure polymerizes from the membrane anchor and ends at the opposite side of the cytoplasmic membrane and because phage sheath contraction is believed to be triggered by baseplate changes resulting from phage fiber attachment to target receptors ^20^, it is possible that sheath contact with the cytoplasmic membrane at the distal end may generate a physical force to the T6SS baseplate, which subsequently causes baseplate conformational change leading to contraction. Although short yet contractible T6SS structures have been found ^34^, cellular geometry makes it difficult to rule out possible membrane contact. However, testing this physical force-driven contraction hypothesis directly poses a technical challenge because it is unfeasible to either remove the cytoplasmic membrane in a living cell or to assemble a functional T6SS *in vitro*.

## Results

### Physical force of membrane contact results in curved structures but does not trigger contraction

Here we designed an alternative strategy by inducing irregular intercellular membrane vesicles in cell wall-deficient cells, as previously reported in *Listeria* L-form cells ^35,36^. We postulate that, if such internal membrane barriers could terminate T6SS sheath growth and trigger contraction, it would strongly support membrane contact as a sufficient trigger (Figure 1A). We found that ampicillin-treated *Vibrio cholerae* spheroplast cells can accumulate internal membrane vesicles (Figure 1B) and decided to use it as a model.

**Figure 1.**
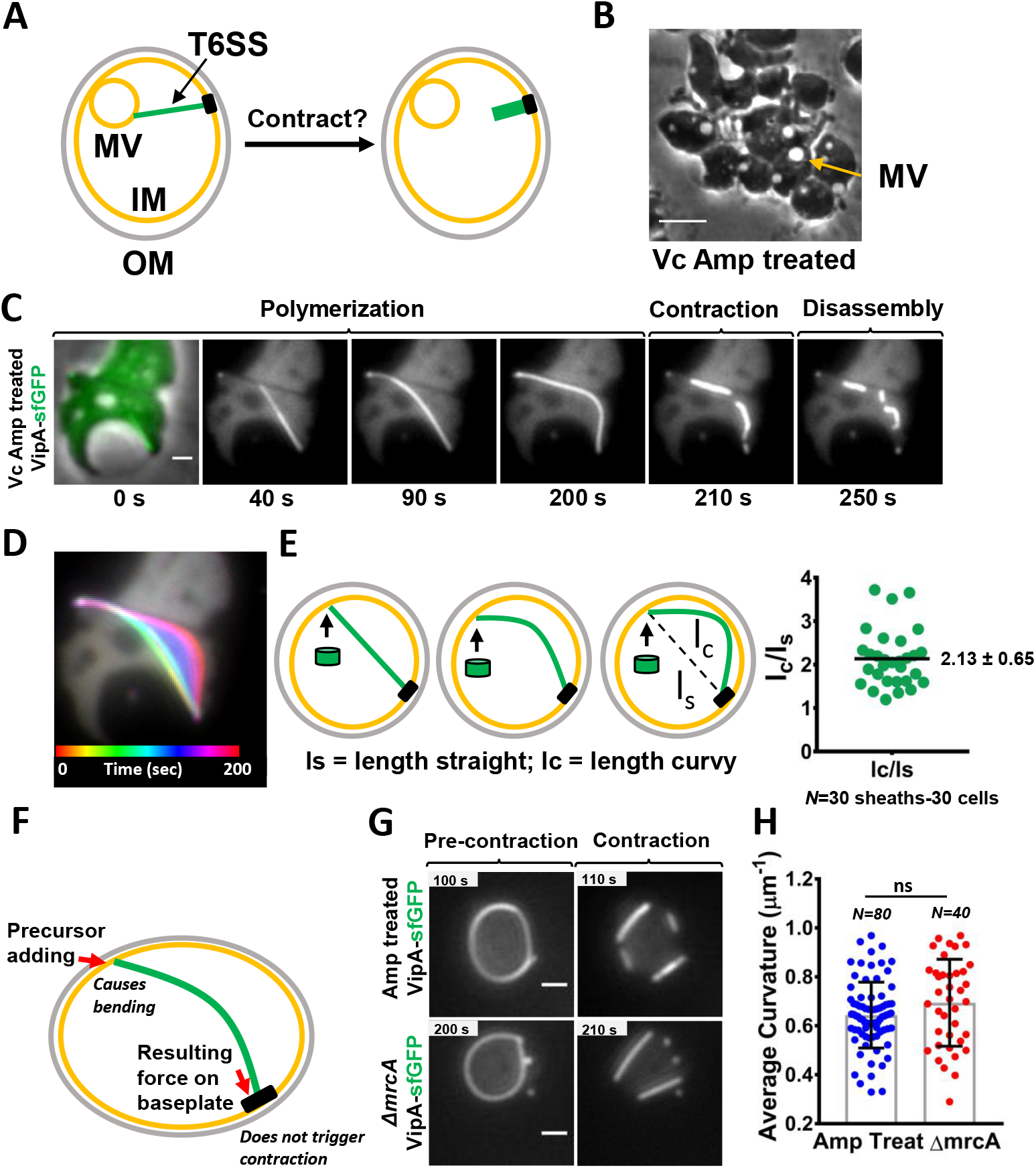
Formation of curved sheaths under membrane vesicle inducing conditions. **A.** Model of internal membrane vesicle (MV) as a trigger for T6SS contraction. IM, inner membrane. OM, outer membrane. **B.** Intracellular membrane vesicles (yellow arrow) in *V. cholerae* (Vc) after 45 min of ampicillin treatment (500 μg/mL) and incubation on agarose pad for 3 h. Scale bar, 5 μm. **C.** Time-lapse showing sheath assembly in the VipA-sfGFP labeled strain treated with ampicillin. Left image is a merge of the phase and GFP (green) channels. Image series correspond to a 5 min time-lapse, 10 sec per frame. Scale bar, 1 μm. **D.** Temporal color coded image corresponds to sheath polymerization (0 to 200 sec) of cell in **C. E.** Model of initially straight sheath (green) followed by curved polymerization. Green cylinders represent sheath subunits. Scatter plot on the right shows the ratio of the total length curvy (1_C_) over the initial/hypothetical straight length (1_S_) for each sheath, and 30 sheaths were measured. **F.** Model shows sheath (in green) bending by continual addition of precursors at the distal end and the resulting force pushing the baseplate (black rectangle). **G.** Pre-contraction and contraction time points of curved sheaths formed by ampicillin-treated (top panel) and *mrcA* deletion (bottom panel) cells labelled with VipA-sfGFP. Full image sequence is shown in Supplementary Figure S1C. Images are representative of at least three biological replicates. Scale bars, 1 μm. **H.** Average curvature of sheaths formed in ampicillin-treated (Mean=0.64 μm^−1^ ± 0.13; n=80, blue dots) and *mrcA* deletion (Mean=0.69 μm^−1^ ± 0.17; n=40, red dots) cells. Scatter plot shows mean value in gray bars±SD (black lines). Significance was calculated by unpaired two-tailed *t*-test. ns, not-significant.

Surprisingly, under vesicle-inducing conditions, we observed highly curved T6SS structures instead of vesicle-triggered early contraction as we initially expected (Figure 1C&D, Supplemental Figures S1A&B, Movie S1). Upon contact, the membrane did not trigger contraction but rather guided a polymerizing straight sheath to bend along and continue to extend. After the sheath’s distal end contacted the membrane, we observed an arching deformation suggesting there is continual addition of VipA/B and Hcp precursors to the growing structure (Figure 1C, D &E). The curved sheaths can still increase their length on average by twofold in comparison with the initial straight length or the linear distance from baseplate to the distal end (Figure 1E). The deformation also suggests that sheath growth generates a bending force that directly presses on the T6SS baseplate-membrane anchor and is counteracted by the physical strength of the cytoplasmic membrane (Figure 1F). Notably, such force is strong enough to bend the sheath-needle double tubular structure but insufficient to trigger contraction (Figure 1F). There are also curved extending sheaths that eventually stopped extending without meeting any physical barriers and in some cases, the polymerizing sheaths fully encircled the whole circumference of the cell (Figure 1G).

To rule out the possibility that the curved extended sheaths are a result of some unknown properties of ampicillin other than targeting cell-wall synthesis, we examined the T6SS formation in the deletion mutant of *mrcA*, encoding the cell-wall synthesis enzyme PBP1a whose absence is known to disrupt the cell wall and causes a spherical shape in *V. cholerae* ^37,38^. We found that curved T6SS sheaths were similarly formed in both, ampicillin-treated cells and the *mrcA* mutant (Figure 1G, Supplemental Figure S1C). We measured the curvature of 120 curved sheaths and found that the average curvature of the flexible sheaths formed in ampicillin-treated cells is 0.64 μm^−1^ (± 0.13; n=80), which is comparable in the *mrcA* mutant (0.69 μm^−1^ ± 0.17; n=40) (Figure 1G). When we measured the curvature at each point along individual sheaths, maximum curvature points reached up to 2.3 μm^−1^ (Mean= 1.22 μm^−1^ ± 0.34; n=120) (Supplemental figure S1D). These results demonstrate the remarkable flexibility of the T6SS double tubular extending sheath/Hcp structure, previously described as a rigid structure, and indicate that contact with the cytoplasmic membrane is not a direct trigger for contraction.

### Determination of curvy limit of extended sheath

Considering the six-start helical strands of the sheath and its inner Hcp tube comprising hundreds of rings of VipA/B and Hcp hexamers, we then asked what might have changed at each sheath ring that collectively account for the observed curvature. For simplicity, we postulate that the bending extending force leads to the stretching of sheath interaction with increased height/rise on one side of each sheath ring when curving (Figure 2A). We define the ratio of height change as a stretching factor (*S*) that is an intrinsic parameter of the sheath-tube structure dependent on the strength of the protomer interactions.

**Figure 2.**
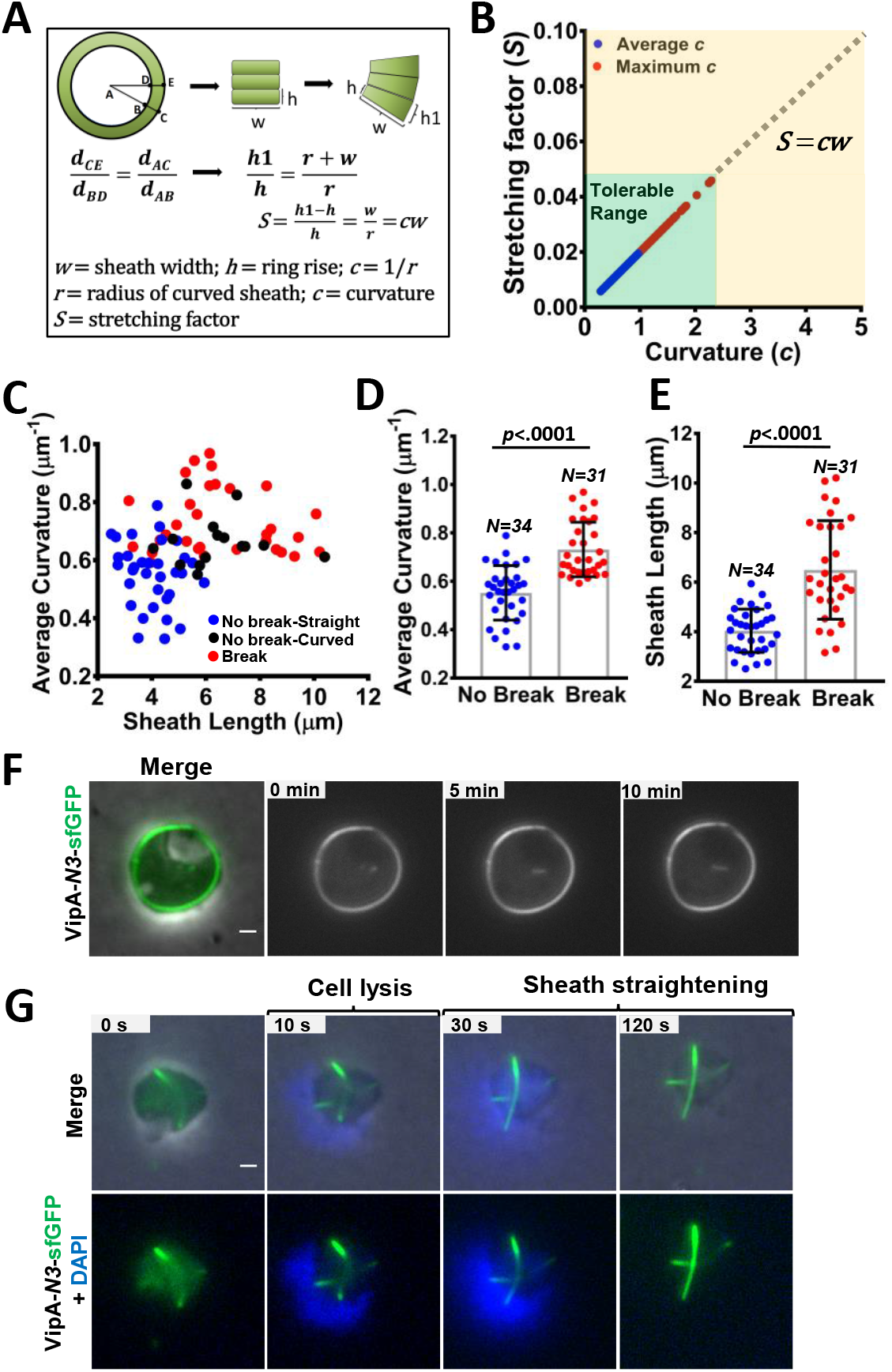
Alleviation of T6SS sheath curvature upon contraction. **A.** Mathematical model to estimate the stretching factor (*S*) of T6SS sheath/Hcp structure. *S* is determined as the ratio of sheath width to the radius (*w/r*) or the product of curvature and sheath width (*c × w*). **B.** Graph shows the tolerable (light green box) and intolerable (yellow area) range of stretching (*S*) (*y*, axis) and curvature (*c*) (*x*, axis) for T6SS sheaths. Plotted values correspond to the average (blue dots) and the maximum (red dots) curvature for each observed curved sheath (n=120). Gray dashed line represents theoretical values. *w*, sheath width. **C.** The scatter plot shows the dispersion of the sheath length and the average curvature (*x, y*) of the three types of contraction events observed in ampicillin-treated cells; normal contraction straightened (blue dots, n=34), normal contraction slightly curved (black dots, n=15) and sheaths that break into pieces after contraction (red dots, n=31). Average curvature **(D)** and sheath length **(E)** of *no break* (blue dots) *and break* (red dots) contraction events are plotted. Scatter plots show mean value in gray bars±SD (black lines). Each data point represents a sheath. Significance was calculated by unpaired two-tailed *t*-test. *p* values are indicated for each graph. Measurements were taken from at least three biological replicates. **F.** Image sequence shows non-contractile curved sheath assembled in the VipA-N3-sfGFP ampicillin-treated strain. Image on the left is a merge of phase and green (GFP) channels. **G.** Non-contractile VipA-N3-sfGFP curved sheath straightening after cell lysis induced by colistin and Triton X. To highlight the lysis event, nucleic acid was stained with DAPI. Top row, merge of phase, GFP (green) and DAPI (blue) channels. Bottom row, green and blue channels only. Full video is included in Movie S3, cell1. Additional example can be found in Supplemental Figure S2D. All scale bars, 1 μm.

Because the width of the sheath (*w = 20 nm*) and rise of each ring (*h = 3.8 nm*) are known in *V. cholerae* ^13^, and the radius for the curved sheaths (*r*) can be measured (Figure 1G), we could estimate the stretching factor (*S*) equal to the ratio of sheath width to the radius (*w/r*) or the product of curvature and sheath width (*c × w*) (Figure 2A). When considering average curvature (*c* ≤ 1 μm^−1^) and assuming that the stretching force is equally distributed along the sheath, we estimated that *S* is less than 2%, and the maximum change of each ring is 0.08 nm and about 0.04% change relative to the sheath width. If local maximum curvature is considered (*c* = 2.27 μm^−1^) (Supplemental Figure S1F), 4% of stretching could be reached at certain points of the sheath, which corresponds to a rise change of 0.16 nm on each affected ring. However, because curvature is measured based on the 2-D projection of the 3-D sheath structure, such maximum curvature is likely an overestimation. By comparison, the rise of sheath decreases from 3.80 nm to 3.78 nm after contraction in *V. cholerae*, and the T6SS extended sheath rise is 3.7 nm in *Myxococcus xanthus* ^13,39^. Importantly, *w* and *S* are intrinsic parameters of the sheath/Hcp structure that ultimately dictate the flexibility potential of the T6SS and define a tolerable range of curvature (Figure 2B). Indeed, we noticed that curvy sheath formation is dependent on the geometry of the cell-sheath contact region. If the contact angle is close to orthogonal, the theoretical curvature at the contacting point will far exceed the maximum tolerable curvature value and thus no curved structures can be formed (Supplemental Figure S1E).

### Curved sheath structure is elastic and restores to straight forms after contraction

Though extended sheaths are curved, contraction reduces the curvature greatly, leading to either straight contracted sheaths (70%, n=34) or much less curved contracted sheaths (30%, n=15) prior to their disassembly (Supplemental Figures S2A,B&C). Interestingly, curved sheaths with a higher curvature value and longer sheaths tended to break at the highest curvy points into straight short pieces due to contraction, prior to disassembly (Figure 2C, D&E, Supplemental Figure S1D). The contracted and break-free sheath pieces were similarly disassembled, suggesting the disassembly process does not require attachment to baseplate (Supplemental Figure S2A). Considering contraction separates at least one end of the sheath from the membrane, the straightening of contracted sheaths suggests that the more compact contracted structures might be less flexible to adopt the curved form and membrane contact is required to confine the sheath into the curvy form.

Next, we asked if the curved extended structures are elastic rather than fixed into a permanent shape. We postulated that removal of membrane constraint would straighten the sheaths. To test this and avoid the complexity of contraction, we used the *vipA-N3* mutant in which the sheath-Hcp tube is locked into the extended form ^40^. The non-contractile extended sheath could assemble curved structures similar to wild type while no contraction was observed (Figure 2F, Movie S2). When cells were treated with detergent TritonX and polymyxin B to induce lysis, the curved structures straightened after escaping from the cells (Figure 2G, Supplemental Figure S2D, Movie S3), indicating that the extended sheath structures are indeed elastic.

### Curved T6SS structures maintain functionality

To test if the curved T6SS sheaths represent functional T6SS structures in delivering effectors, we used a sensitive T6SS complement assay using the T6SS null *hcp1/2* and *vgrG2* mutants as reporter ^34^ The reporter strains cannot assemble T6SS sheath/Hcp structure due to the lack of key structural components unless the missing components are provided through delivery by a donor cell ^34^. Indeed, under the curvy-sheath inducing conditions, we readily detected sheath formation in both reporter strains while none detected when the donor was the T6SS mutant (Figure 3A, Supplemental Figures S3A&B). More sheaths were observed in the *hcp* mutant while longer sheaths were observed in the *vgrG2* mutant (Figure 3B), reflecting the fact that one VgrG trimer and a stack of Hcp hexameric rings are needed to assemble a functional extended sheath-Hcp structure.

**Figure 3.**
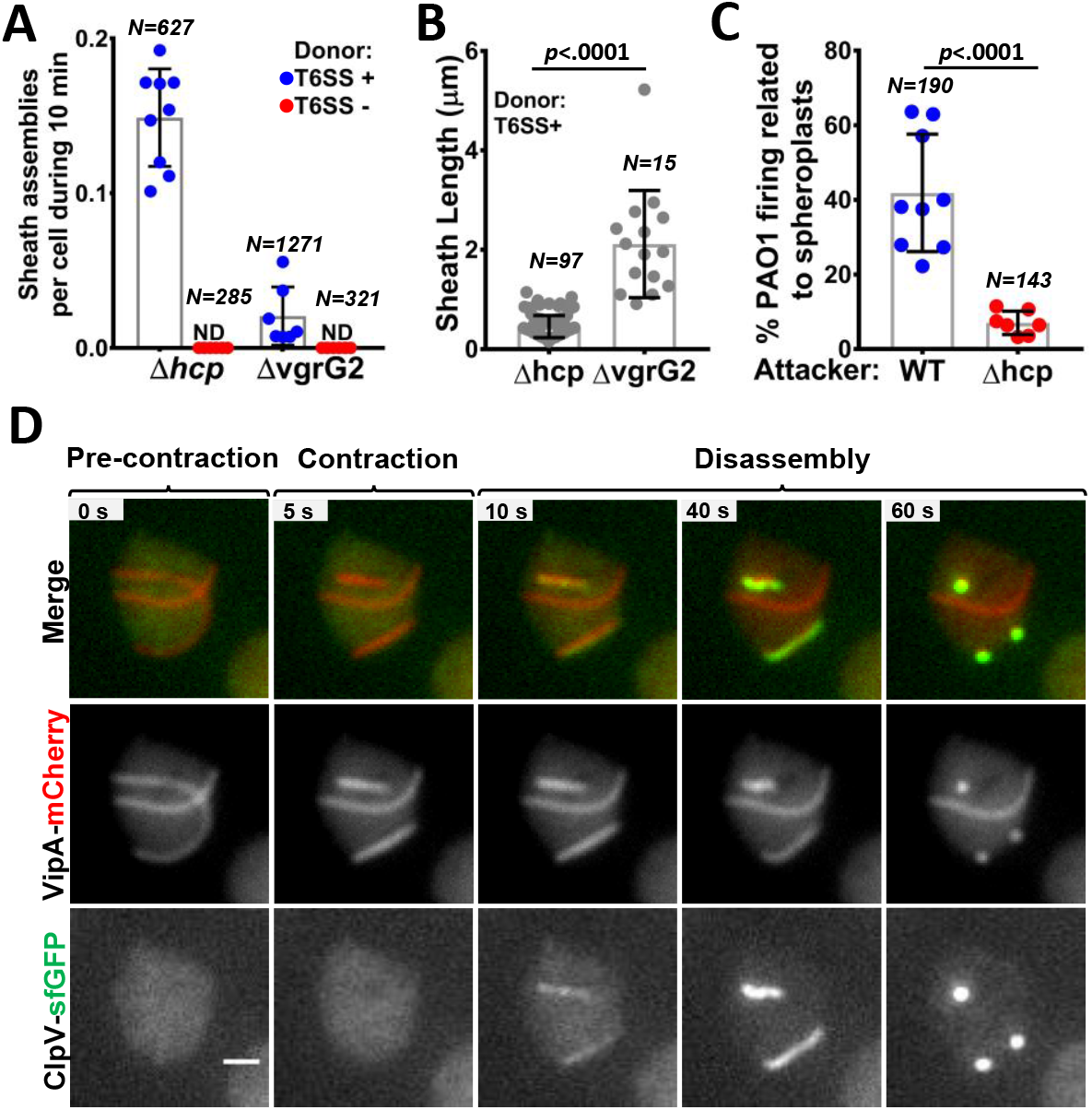
T6SS remains functional under curved sheaths condition. **A.** The number of sheaths assembled in the VipA-sfGFP recipient strains (Δ*hcp* and Δ*vgrG2*) during 10 minutes in co-incubation with either, the wild type (WT) VipA-mCherry (donor T6SS+, blue dots) or unlabelled Δ*vipB* strain (donor T6SS-, red dots) is shown. Both recipient and donor cells were treated with ampicillin and mixed in a 3:1 ratio (donor to recipient) for imaging. Three videos from each of the three biological replicates were analyzed for each condition. The number of cells counted (*N*) is indicated at the top. ND, not detected. **B.** Sheath length (μm) was measured for each sheath formed in both recipient strains, the VipA-sfGFP Δ*hcp* and the VipA-sfGFP Δ*vgrG2*, co-incubated with the T6SS+ strain. The number of sheaths observed (*N*) is indicated at the top. **C.** Percentage of foci assembled by *P. aeruginosa* PAO1-TssB1-sfGFP surrounding either, WT (blue dots) or Δ*hcp* (red dots)-VipA-mCherry *V. cholerae* spheroplasts, over total number of observed foci during 15 minutes. Each data point represents a video from three biological replicates. Total number of foci observed (*N*) is indicated at the top. **D.** Image sequence of a double labelled VipA-mCherry ClpV-sfGFP cell showing curved sheath pre-contraction, contraction and subsequent disassembly by ClpV. Frames on the top row correspond to a merge of mCherry (red) and GFP (green) fluorescence channels. Middle and bottom rows show VipA-mCherry and ClpV-sfGFP signals only, respectively in grayscale. Image sequences for panels A, B&C and, an additional example for panel D are included in Supplemental Figure S3. Scale bar, 1 μm. In **A, B** and **C**, scatter plots show mean value in gray bars±SD (black lines). Significance was calculated by unpaired two-tailed *t*-test. *p* values are indicated for each graph.

The H1-T6SS in *Pseudomonas aeruginosa* PAO1 can respond to external T6SS attacks and thus can be used as a reporter for T6SS delivery as well ^41^. Indeed, under the curvy-sheath inducing condition, significantly more T6SS activities in PAO1 were observed when PAO1 cells were neighboring wild type *V. cholerae* than when surrounding the *hcp* mutant (Figure 3C, Supplemental Figure S3C).

Next we tested whether the curved sheath could contract normally and be disassembled by the ATPase ClpV. The otherwise buried VipB N-terminal domain is exposed only after sheath contraction and specifically recognized by ClpV that disassembles the contracted sheath ^21–24^. The curved extended sheaths were not recognized by ClpV but contraction quickly led to ClpV coating of the sheath, indicating the VipB N-terminus is inaccessible in the curved extended sheath the same as in the straight extended sheath (Figure 3D, Supplemental Figure S3D, Movie S4). These results collectively indicate that the curved extended sheath-tube structures remain functional.

### Stalling is an unstable intermediate stage between extension and contraction

Imaging analysis shows that in both rod (n=150) and spheroplast (n=80) cells, there is a stalling stage (~ 50 seconds, median value) during which the extended sheath maintains unchanged (Figure 4A, Supplemental Figure S4A). After the stalling, the extended sheath could continue to extend or undergo contraction. T6SS sheath assembly is also strictly dependent on, and proportional to, the availability of Hcp under curvy sheath inducing conditions, indicating that the observed curvy structures are not due to abnormal assembly of empty sheath (Figure 4B, Supplemental Figure S4D). Interestingly, despite the length difference, the number of sheaths per cell remains comparable under different Hcp inducing levels (Figure 4C), suggesting that the T6SS assemblies initiating from different baseplate complexes could compete for limited amounts of Hcp. In addition, VipA-sfGFP precursor levels fluctuated in inverse correlation with the total length of polymerized sheath, indicating an intracellular recycling balance between free and polymerized sheath components (Figure 4D, Supplemental Figures S4B&C).

**Figure 4.**
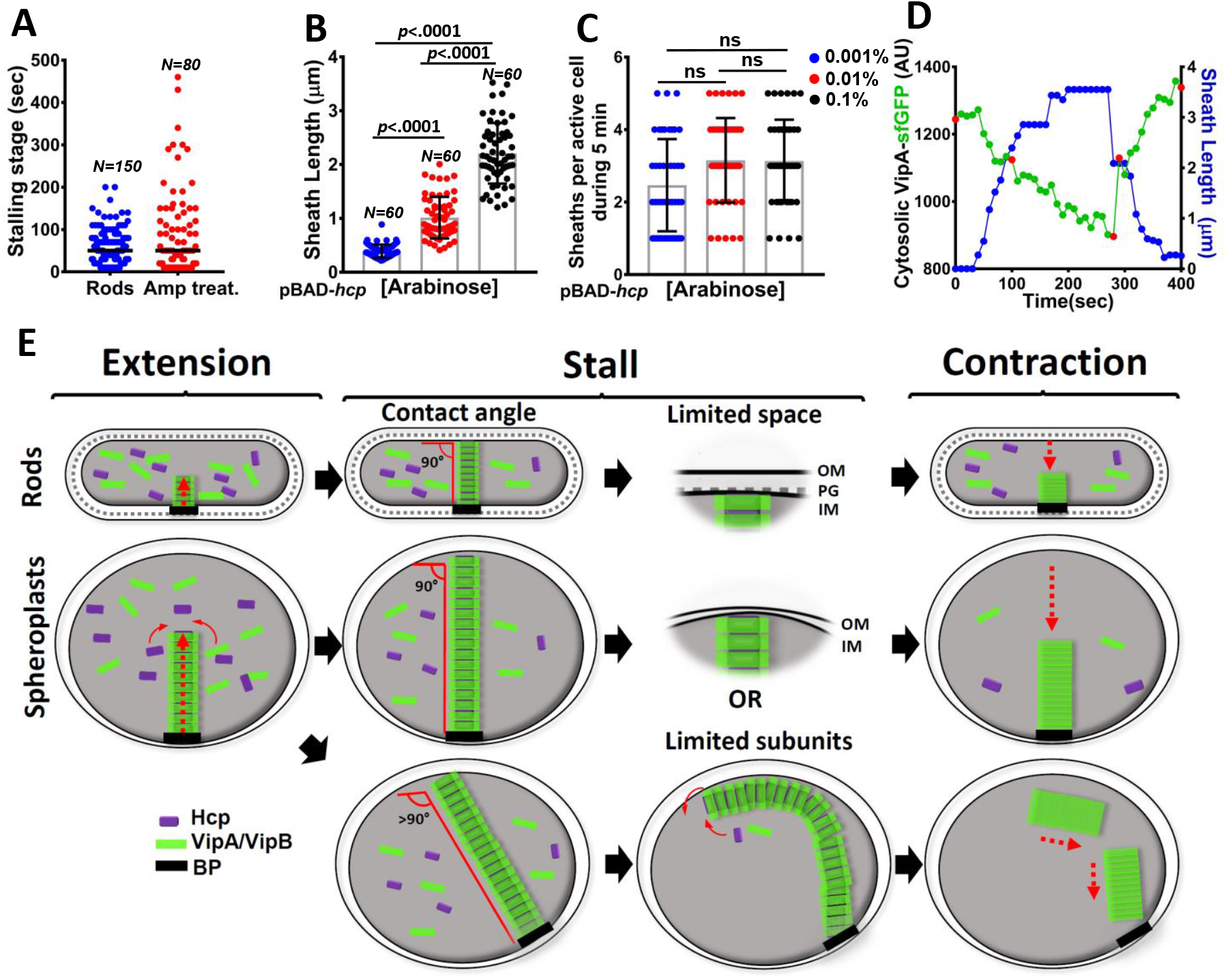
Model of the T6SS sheath/Hcp tube extension and contraction. **A.** Stalling stage, represents the time between full extension and contraction in rod cells (blue dots, n=150) and spheroplasts (red dots, n=80). The median value (= 50 sec) is indicated by a black line. **B** and **C** show the length (n=60 sheaths per condition) and the number (n=40 cells per condition) of sheath assembled upon *hcp* induction in spheroplast cells, respectively. The VipA-sfGFP *hcp* deletion mutant carrying pBAD18-*hcp1* plasmid was induced under increasing concentrations of L-arabinose, 0.001% (blue dots), 0.01% (red dots) and 0.1% (black dots). Scatter plots show mean value in gray bars±SD (black lines). **D.** The fluorescence intensity of VipA-sfGFP in the cytosol (excluding sheath) (green, left y axis) and the length of the polymerizing sheath (blue, right y axis) at each time point (x, axis) of a 5 min time-lapse video (10 sec per frame) was measured in a representative spheroplast. The microscopy images corresponding to the time points indicated in red are shown in Supplemental Figure S4B. Additional example is shown in Supplemental Figure S4C. **E.** ESCAPE (extension-stall-contraction and precursor equilibrium) model. Extension, stall and contraction stages are schemed for rods (top) and spheroplasts (bottom). Significance was calculated by One-way ANOVA with multiple comparison using Tuckey post hoc test in **B** and **C.** *p* values are indicated for each graph. ns, not-significant. All measurements were taken from at least three biological replicates.

### ESCAPE (extension-stall-contraction and precursor equilibrium) model for sheath-tube extension and contraction

Based on these observations, we propose that the T6SS sheath-tube length and contraction are dependent on an equilibrium of stabilizing precursor addition and destabilizing tendency for contraction (Figure 4E). Analogous to an enzymatic reaction, precursors of VipA/B sheath and Hcp tube tend to polymerize to form the extended sheath-tube structure. However, polymerization enters a stalling stage when no new precursor is added, either due to depletion of precursor or limitation of physical space (most likely in rod cells). When precursors can be recruited again, either by recycling of VipA/B sheath subunits and *de novo* synthesis of Hcp, or the distal end is freed due to membrane flexibility, the stalling structure restarts extension. Otherwise contraction occurs and ejects the inner Hcp tube and effectors outward.

## Discussion

Contractile needle-like cell-puncturing devices are often involved in delivering toxins into target cells. The T6SS is mechanistically and evolutionarily related to R-type pyocins ^42,43^, *Photorhabdus* virulence cassettes (PVC) ^44^, *Serratia* anti-feed prophage ^45^, and contractile phage tail ^46,47^. Their built-in strength for puncturing and distinct concentric double tubes implicate the structural rigidity. The sheath is assembled into a high energy extended state and contraction completes almost instantly (less than 2 milliseconds for T6SS) ^10^, which enables a powerful penetration to breach the target cell. Here we report that the T6SS structure is not rigid but rather strikingly flexible and elastic. The double tubular sheath-tube structure could accommodate physical force to deform while maintaining connectivity and functionality. It is generally believed that the baseplate of evolutionarily related contractile phages switches to contraction by sensing physical changes resulting from the attachment of tail fibers to receptors ^20,48,49^. By contrast, we demonstrate that the pressing force, resulting from sheath-membrane contact and acting on the T6SS membrane-baseplate complex, could cause straight to curvy deformation but could not trigger contraction (Figures 1C&D).

Based on our results and previous findings that T6SS sheath length correlates with Hcp abundance and physical space ^10,34^, we propose a precursor-equilibrium-mediated stability model (Figure 4E). The effect of precursor abundance on sheath stability could be direct and/or indirect. In the former scenario, precursor recruitment could directly transmit a stabilizing signal from the distal end to the baseplate of an extending sheath. Such stabilization may involve a conserved T6SS protein TssA that directly interacts with the sheath-tube precursors Hcp and VipB ^50,51^. It has been proposed that a dodecameric TssA complex functions as a cap at the sheath distal end to recruit sheath-tube precursors and stabilize the structure ^10,51^, while another model suggests that TssA remains attached to the baseplate ^50^. Notably, there are two distantly related TssA homologs in *V. cholerae*. Deletion of *tssA1* but not *tssA2* results in abolished T6SS secretion ^52^. However, the exact functions of these two TssAs remain elusive and warrant further studies. Alternatively, precursor might stabilize the extended sheath indirectly by inactivating some unknown contraction-triggering proteins or preventing their interaction with the extending sheath.

Why are the T6SS structures curved and why are they not seen in normal rod cells? All sheath-tube structures are straight while extending until their distal ends touch the membrane. Continual addition of precursors at the distal end would generate a conflict between the tendency to remain straight and the physical confinement of the membrane. Such conflict results in a mechanical bending force exerted on the sheath from the membrane contacting points, mainly the membrane anchor and the distal end (as seen in Figure 1c). The resulting stress could force a range of angle changes at the membrane contacting points in spheroplast cells that lack a rigid cell wall. Such angle change is key to accommodate the continual addition of precursors but is much less likely to occur in rod cells considering that the T6SS essential TssJLM membrane complex spans across the two membranes and the rigid cell wall. We also found straight sheaths at early time points but curvy sheaths at later time points after ampicillin treatment, further supporting that cell wall inhibits curvy sheath formation. Notably, a previous report using similar ampicillin-treatment in *V. cholerae* found long but straight sheath when imaged at earlier time points (40 min) ^10^. It is also possible that some unknown factors controlling the sheath shape are depleted or enriched at later time points in cell wall deficient cells. Such factors may be identified using genetic screening of random transposon mutants combined with high throughput imaging tools.

The six-start helical structure of both the inner and outer tubes ^13^ likely provide the structural basis for the stretching, curvy formation and elasticity. Using time-lapse fluorescence microscopy, we could monitor the dynamic assembly of curved extended sheath and provide a theoretical estimate on the stretching potential of the T6SS structure. The observed flexible T6SS structures are phenotypically reminiscent of the long and flexible tails of non-contractile phages^53^, suggesting a common ancestry.

Considering the relatedness of all known contractile double-tubular structures, the remarkable flexibility of T6SS sheath-tube might represent a general nature of elasticity among similar contractile injection systems.

## Materials and Methods

### Bacterial strains and growth conditions

All strains and plasmids used in this study are listed in Tables S1 and S2, respectively. Cells were grown aerobically at 37 °C in LB (1% [wt/vol] tryptone, 0.5% [wt/vol] yeast extract, 0.5% [wt/vol] NaCl). Antibiotics and inducers were used at the following concentrations: chloramphenicol 25 μg/mL (for *E. coli*) and 2.5 μg/mL (for *V. cholerae*), 20 μg/mL gentamycin, 100 μg/mL streptomycin and 25 μg/mL irgasan. Gene expression vectors and precise knockout mutants were constructed as previously described ^54^. All constructs were verified by sequencing. All primers used in this study are provided in Table S3.

### Induction of Membrane Vesicle (MV) formation by ampicillin treatment

Overnight cultures were diluted 1/100 in 2 mL of fresh LB and grown aerobically at 37 °C to an OD_600_ of 0.6-0.8. Ampicillin was added to the cultures at 500 μg/mL and further incubated for 45 min. Culture was centrifuged and pellet was resuspended in 0.5X PBS buffer. Cells were spotted onto 1% agarose-pads supplemented with diluted LB (1:5) and PBS buffer (1:2), and ampicillin (10 μg/mL). Spotted microscopy slides were incubated at 37 °C and imaged at different time points.

#### *V. cholerae AmrcA* spheroplasts

Overnight culture of the Δ*mrcA* strain was centrifuged and the pellet was resuspended in fresh LB. After incubation at 37°C for 1 h, cells were concentrated to OD600 =10 and spotted onto 1% agarose-0.5X PBS pads supplemented with diluted LB (1:5). The spotted microscopy slides were incubated at 37 C prior to imaging.

#### Cell lysis induction

*V. cholerae* VipA-N3-sfGFP cells were grown to OD_600_=0.6-0.8 and treated with 500 μg/mL ampicillin for 45 min. Cells were then collected and spotted onto an agarose pad under membrane vesicle inducing conditions, as explained before. After incubation at 37 □C of the spotted microscope slides, the coverslip was carefully removed and 1 μl of 80 μg/mL colistin-0.1% Triton X plus 1 μl of 10 μg/mL DAPI were deposited on top of the agarose pad. A new coverslip was added and cell lysis was quickly monitored by time-lapse fluorescence microscopy at a frame rate of 10 seconds per frame until all observed cells were lysed (approximately 30 minutes).

#### Functionality assays under curvy structure conditions

For the T6SS interbacterial complementation assay, Δ*hcp1/2* VipA-sfGFP and Δ*vgrG2* VipA-sfGFP strains were used as recipient cells. Wild type VipA-mCherry (T6SS +) or Δ*vipB* (T6SS -) were used as donor strains. Both, donor and recipient cells were treated similarly with 500 μg/mL of ampicillin, harvested and then mixed to a 1:3 ratio (recipient to donor) for imaging. Sheath formation in the recipient strains was monitored using time-lapse fluorescence microscopy at a frame rate of 10 seconds per frame for 10 minutes.

For the *P. aeruginosa* PAO1 reporter assay, *V. cholerae* wild-type (T6SS +) and Δ*hcp1/2* (T6SS -) VipA-mCherry strains were grown to exponential phase and treated with ampicillin. Cells were spotted onto 1% agarose-0.5X PBS pads supplemented with LB and ampicillin and incubated to induce curvy sheaths formation. The glass coverslip was carefully removed and, exponentially growing PAO1 TssB-sfGFP rod cells were spotted on top of the *V. cholerae* containing-agarose pads. A new glass coverslip was added and cells were immediately imaged every 10 sec using time-lapse fluorescent microscopy.

To image ClpV-mediated disassembly, *V. cholerae* VipA-mCherry ClpV-sfGFP cells were treated as explained before for curved sheaths formation and visualized after incubation at 37 C on the agarose pad. Cells were imaged by time-lapse fluorescence microscopy for 8 minutes at a frame rate of 5.5 sec per frame.

#### Hcp complementation assay

Exponentially growing cells (0D_600_=0.6-0.8) of *V. cholerae* Δ*hcp1/2* VipA-sfGFP strain either, empty or carrying a pBAD18-*hcp1* plasmid were treated with 500 μg/mL of ampicillin for 45 min and spotted onto agarose pads. To induce Hcp expression, different concentrations (0.001%, 0.01% or 0.1% [wt/vol]) of L-arabinose were added to the agarose pads. Cells were imaged after 2 h incubation at 37□C.

#### Image acquisition

All images were acquired by a Nikon Ti-E inverted microscope with a Perfect Focus System (PFS) and a CFI Plan Apochromat Lambda 100X oil objective lens. Different fluorescence signals were filtered and excited using Intensilight C-HGFIE (Nikon), ET-GFP (Chroma 49002), and ET-mCherry (Chroma 49008) filter sets. Images were recorded by an ANDOR Clara camera (DR 328G-C01-SIL), pixel size 60 nm and, NIS-Elements AR 4.40 software.

#### Image analysis

Fiji was used for all image analysis ^55^. To correct for photobleaching, fluorescence intensity of the acquired time lapse videos was normalized to the same mean intensity for each image in a time series as described previously ^41^. The plugin “Temporal-Color Code” for Fiji was used to visualize the polymerization trajectory of the curved sheath in Figure 1D. Sheath length of straight and curved sheaths was measured using straight and freehand line tools, respectively. To calculate sheath contact angle with the cell membrane, the angle tool for Fiji was used. The curvature analysis was performed using the plugin “Kappa” for Fiji. The curvature is defined as the reciprocal of the radius (*r*^−1^). For sheaths with negative curvature points, the absolute values of curvature were used to build graphs and calculate mean values. Five points were used to define each curved sheath and the scale was set to 0.064 μm/pixel. We measured 120 fully extended curved sheaths (80 from ampicillin treated and 40 from Δ*mrcA* spheroplasts). To quantify cytosolic VipA-sfGFP fluorescence intensity, the average mean gray value of five circles (0.13×0.13 μm) distributed in the cytosol of the cell minus average fluorescence background outside the cell was calculated at each time point. “KymoResliceWide” plugin for Fiji was used to generate kymographs. Freehand line tool was used to define the sheath area to generate the kymographs.

#### Quantification and statistical analysis

The unpaired two-tailed t-test as well as one-way ANOVA with multiple comparisons and Tuckey post hoc test were performed using GraphPad Prism version 7.04. Number of cells analyzed as well as significance of each comparison are indicated in figure and figure legends. All experiments were performed with at least three biological replicates. Unless indicated differently, values are expressed as mean±SD.

## Acknowledgments

This work was supported by grants from the Canadian Institutes of Health Research and the Natural Sciences and Engineering Research Council (NSERC) to T.G.D. T.G.D. is supported by a Canada Research Chair award. We thank members of the Dong lab for providing reagents, general support and critical reading of the manuscript.

## Contributions

T.G.D. conceived the project. M.S.S, X.L. and T.G.D. designed the experiments. M.S.S., X.L., M.W., and S.H. performed the experiments. M.S.S. and T.G.D. prepared the manuscript.

## Supplemental figure legends

**Figure S1.**
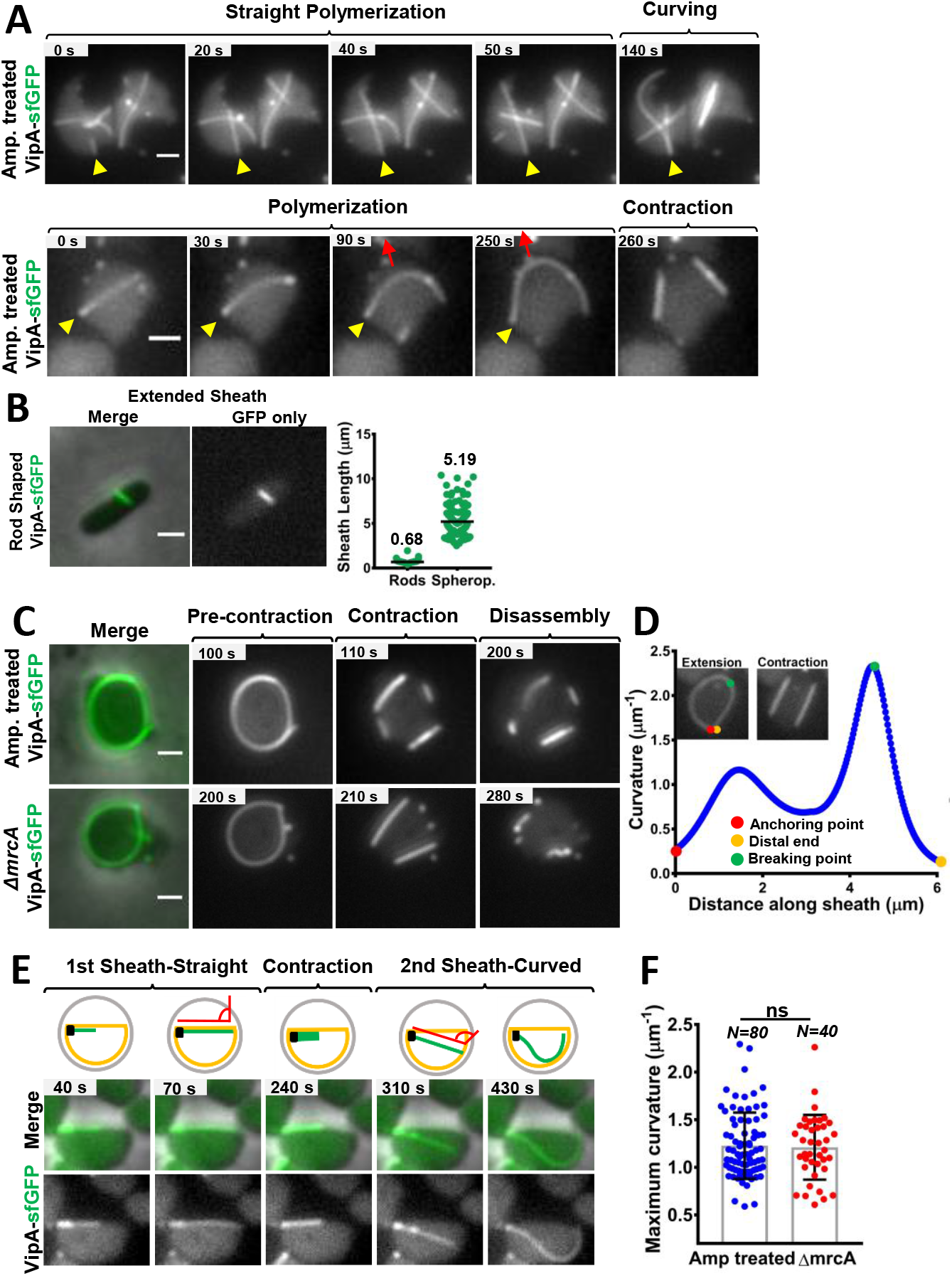
Arching deformation of the T6SS sheath. Related to Figure 1. **A.** Two examples of sheath polymerization are shown. Image sequences correspond to VipA-sfGFP labelled spheroplasts under vesicle inducing conditions. The GFP (green) fluorescence channel is shown in grayscale. Yellow arrows indicate anchoring point. Red arrows in the bottom row indicate direction of sheath arching deformation. **B.** An image of a sheath assembled in a VipA-sfGFP rod cell is shown in a merge of phase and GFP channels (left) and GFP channel only (right). The graph on the right shows the sheath length distribution in rod cells (mean=0.68 μm), and spheroplasts (mean=5.19 μm). **C.** Image sequences, related to Figure 1G, show curved sheaths formed by the VipA-sfGFP labelled ampicillin-treated (top panel) and *mrcA* deletion (bottom panel) cells. **D.** Sheath curvature (y, axes) was measured at each point along the sheath (x, axes) in VipA-sfGFP labelled spheroplasts treated with 500 μg/mL ampicillin. 3 x 3 μm insets show the extended and contracted state of the sheath. The sheath anchoring, distal end and breaking point are shown in red, yellow and green dots, respectively. **E.** Contacting angle of sheath with the cell membrane. A 90° sheath assembles straight (1^st^ sheath), followed by 110° sheath that curves (2^nd^ sheath). Microscopy image sequence is shown as a merge of phase and green channels (middle row) and green channel only (grayscale, bottom row). The sequence is schematized in the top panel. **F.** The curvature at each point along the sheath was measured for curved sheaths formed in ampicillin-treated (Mean=1.22 μm^−1^ ± 0.34; n=80, blue dots) and *mrcA* deletion (Mean=1.21 μm^−1^ ± 0.33; n=40, red dots) cells, as shown in C. The maximum curvature value for each needle is plotted. Scatter plot shows mean value in gray bars±SD (black lines). Significance was calculated by unpaired two-tailed *t*-test. *p* values are indicated for each graph. All scale bars, 1 μm.

**Figure S2.**
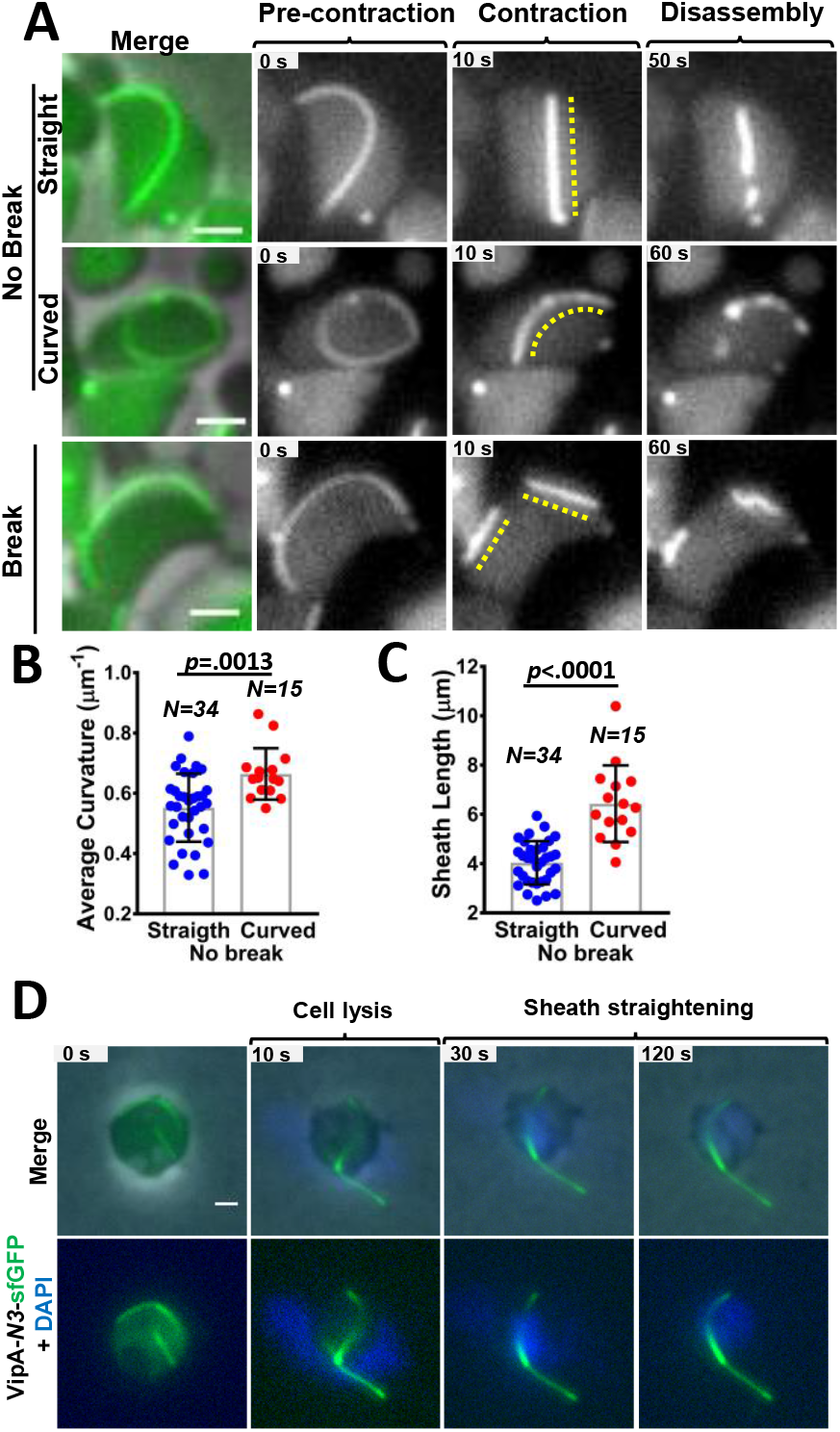
Different contraction events in curved sheaths. Related to Figure 2. **A.** Image sequences show sheaths formed in ampicillin-treated cells that contract normally (*No break*) and straighten after contraction (Top row), contract normally (*No break*) but remain slightly curved (Middle row) and that break into pieces after contraction (Bottom row). Yellow dash line indicates the position of the contracted sheath. Left image is a merge of the phase and GFP (green) channels. Images were taken from a 5 min time-lapse, 10 sec per frame. Average curvature (**B**) and curve length (**C**) of normally contracting sheath (*No break*) formed in ampicillin-treated cells that, either straighten (n=34, blue dots) or remain slightly curved (n=15, red dots) after contraction, are shown. Scatter plots show mean value in gray bars±SD (black lines). Each data point represents a sheath. Significance was calculated by unpaired two-tailed t-test. *p* values are indicated for each graph. ns, not-significant. ns, not-significant. **D.** Additional example of straightening after cell lysis, related to figure 2G, induced by 80 μg/mL colistin-0.1% Triton X. Top row, merge of phase, green (VipA-sfGFP) and blue (DAPI) channels. Bottom row, green and blue channels only. Full video is included in Movie S3, cell 2. Scale bar, 1 μm.

**Figure S3.**
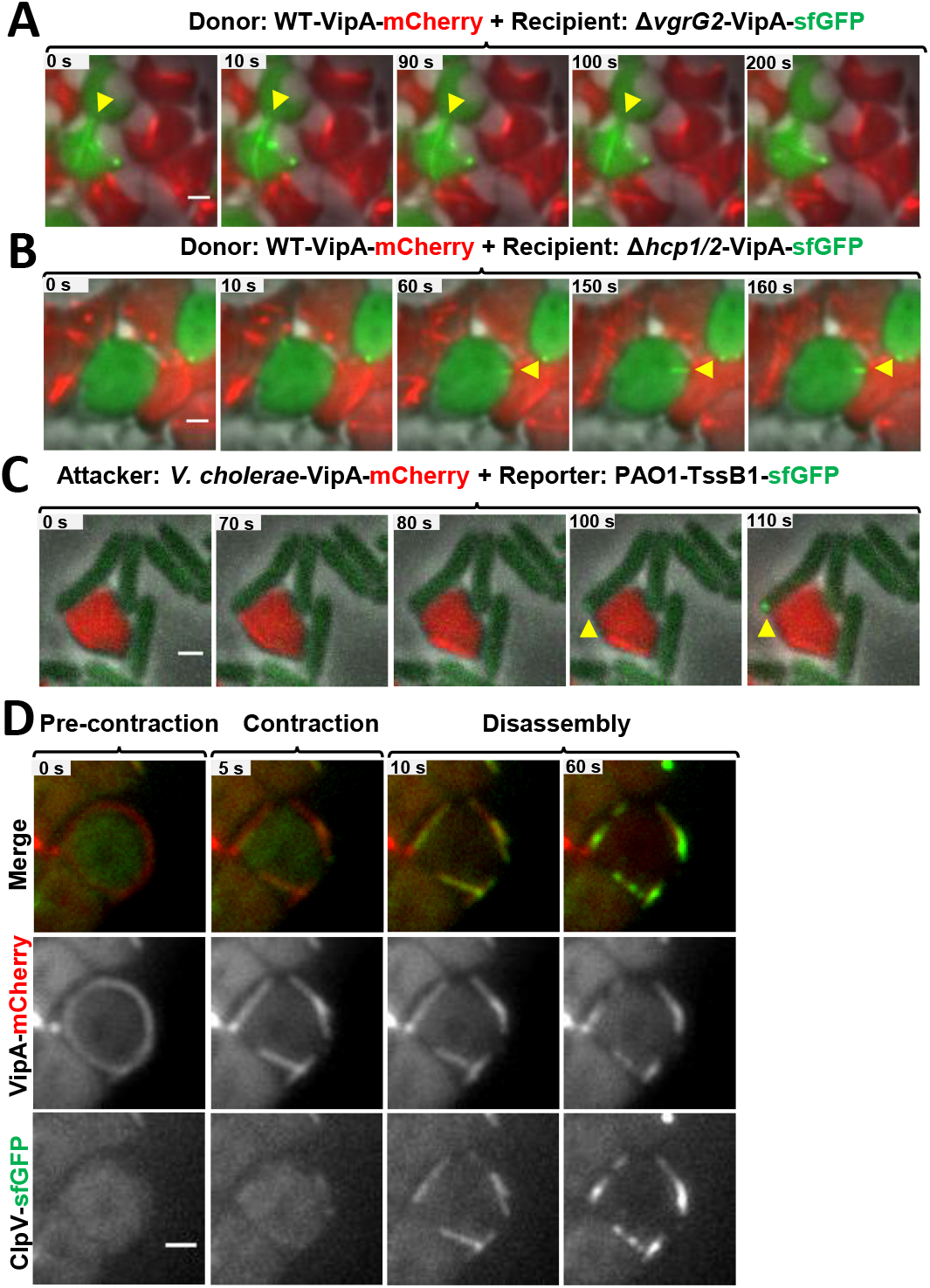
Curved T6SS structures deliver effectors and provoke PAO1. Related to Figure 3. Image sequences show representative examples of sheath formation in the recipient cells, Δ*vgrG2*-VipA-sfGFP (**A**) and *Δhcp*1/2-VipA-sfGFP (**B**) co-incubated with the donor strain, T6SS+ VipA-mCherry. Both recipient and donor cells were treated with 500 μg/mL ampicillin for 45 min, mixed in a 3:1 ratio (donor to recipient) and incubated onto an agarose pad for 3 h before imaging. Each image is a merge of phase, mCherry (red) and GFP (green) channels. Sheath formation in the recipient cells is indicated by yellow arrows. **C.** A representative example of a focus formed (yellow arrows at 100 and 110 seconds time points) in the reporter strain *P. aeruginosa* PAO1-TssB-sfGFP after sheath contraction in the attacker cell, *V. cholerae* T6SS+ VipA-mCherry. **D.** Additional example of a curved circular sheath assembled in a VipA-mCherry-ClpV-sfGFP ampicillin-treated cell. Pre-contraction, contraction and disassembly by ClpV are shown. Top row corresponds to a merge of mCherry (red) and GFP (green) fluorescence channels. Middle and bottom rows show VipA-mCherry and ClpV-sfGFP signal only, respectively in grayscale. 5 minutes time-lapse, frame rate, 5.5 sec per frame. All scale bars, 1 μm.

**Figure S4.**
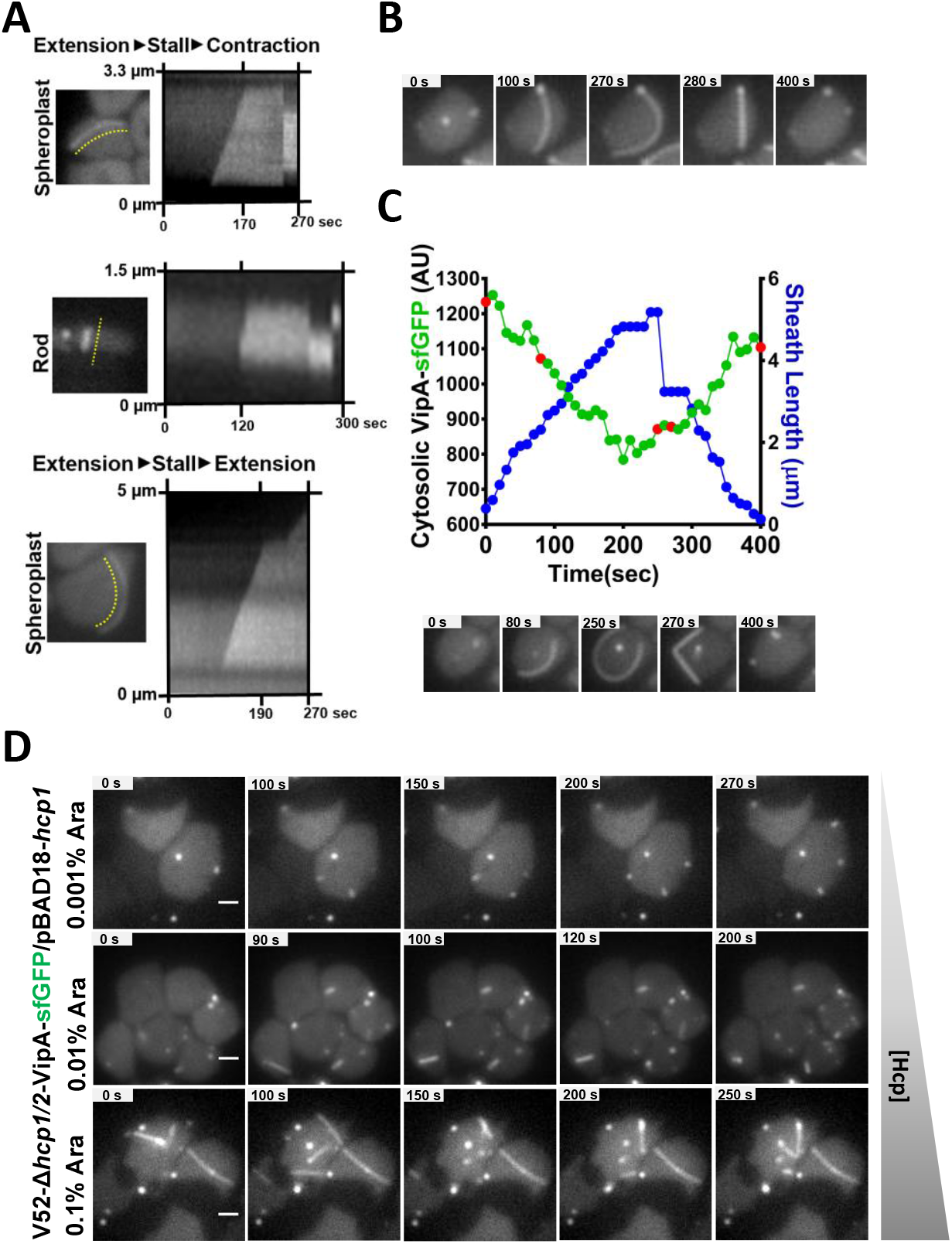
Sheath/Hcp tube dynamics related to precursor availability. Related to Figure 4. **A.** Kymographs correspond to the area indicated by yellow dash lines. 5 min time-lapse videos of 5.5 sec (for spheroplasts) and 10 sec (for rod) interval per frame were used to build the kymographs. Top and middle rows show examples of sheath stall stage before contraction in spheroplast (=70 sec) and rod (= 90 sec) cells. Bottom row shows example of sheath stall stage (= 45 sec) in spheroplast before restarting polymerization. **B.** Image sequence corresponding to Figure 4D. **C.** Additional example of fluorescence intensity analysis of VipA-sfGFP in the cytosol (green, left *y* axis) and the length of the polymerizing sheath (blue, right y axis). 5 min time-lapse, 10 sec per frame. Microscopy images of the analyzed cell is shown on the bottom row corresponding to the time points indicated in red. **D.** Image sequences show different lengths of sheath assemblies during 5 minutes time-lapse videos in the VipA-sfGFP *hcp* deletion mutant carrying the pBAD18-*hcp1* plasmid under increasing (top to bottom) concentrations of inducer (0.001%, 0.01% and 0.1% L-arabinose). GFP channel in grayscale is shown for all images. Scale bars, 1 μm.

## Supplemental Movie legends

**Movie 1. Curved sheath formation in *V. cholerae* spheroplasts.** Related to Figure 1 and Supplemental Figure S1. VipA-sfGFP labelled cells were treated with 500 μg/mL ampicillin for 45 min and incubated on agarose pad before imaging. Two representative fields of view (FOV) of 40 x 40 μm acquired during 15 min time-lapse with a 10 sec frame rate are shown. Video plays at a rate of 10 frames per second. Left field shows a merge between phase (pseudo-colored as blue) and GFP fluorescence channels; right field shows only GFP fluorescence channel in grayscale.

**Movie 2. Non-contractile VipA-N3 curved sheaths.** Related to Figure 2F. The VipA-N3-sfGFP strain was treated with ampicillin for 45 min to induce spheroplast formation. Cells were imaged after 3 h incubation on agarose pad. Three representative videos acquired during 10 min time-lapse with a 10 sec frame rate are shown. Field of view (FOV) is 40 x 40 μm. Video plays at a rate of 10 frames per second. Left field shows a merge between phase and GFP fluorescence channels; right field shows only GFP fluorescence channel in grayscale.

**Movie 3. Cell lysis and sheath straightening.** Related to Figure 2G and Supplemental Figure S2D. The VipA-N3-sfGFP strain was treated with ampicillin and incubated on agarose pad for 3 h. Cell lysis was induced by addition of 80 μg/mL colistin-0.1% Triton X to the agarose pad. To stain nucleic acid, 10 μg/mL DAPI were added. Two representative cell lysis events are shown. Field of view is 10 x 10 μm. Image series were taken during 8 min time-lapse at a 10 sec frame rate. Video plays at a rate of 10 frames per second. Left field shows a merge of phase, GFP (green) and DAPI (blue) fluorescence channels. On the right, field shows GFP and DAPI fluorescence channels only.

**Movie 4. Curved sheaths are disassembled by ClpV upon contraction.** Related to Figure 3D and Supplemental Figure S3D The VipA-mCherry-ClpV-sfGFP double labelled strain was used to induce spheroplasts by ampicillin-treatment. Image series were taken after 3 h incubation on an agarose pad. Three representative fields of view (FOV) acquired during 5 min time-lapse with a 5.5 sec frame rate are shown. Field of view is 40 x 40 μm. Video plays at a rate of 10 frames per second. Left field shows a merge of phase channel, mCherry (red) and GFP (green) fluorescence channels; middle field shows only mCherry channel in gray scale and right field shows only GFP fluorescence channel in gray scale.

